# Delayed dynamics of migratory response to CTLA-4 blockade reveals a mechanistic view on potential T cells reinvigoration following immune checkpoint blockade

**DOI:** 10.1101/2022.04.03.485914

**Authors:** Fateme Safaeifard, Seyed Peyman Shariatpanahi, Bahram Golieai, Amir R. Aref, Mohammad-Hadi Foroughmand-Araabi, Sama Goliaei, Curzio Rüegg

## Abstract

Cytotoxic T-lymphocyte-associated antigen 4 (CTLA-4) and programmed cell death protein 1 (PD-1) receptors, two clinically relevant targets for immunotherapy of cancer, are negative regulators of in immune cell activation and migration. However, optimizing therapeutic outcomes still requires fundamental research to reach a comprehensive insight into the coherent function of immune regulators.

Here, we investigated the statistical dynamics of T cells migration as a measure of the functional response to these pathways in an experimental setup of immune checkpoint blockade. For this purpose, we used a previously developed 3-dimensional organotypic culture of patient-derived tumor spheroids.

Experiment-based dynamical modeling remarked distinct characteristics of the receptors regulation followed through with the modification of their proportions in the immune modulation. We demonstrated that time-delayed kinetics of PD-1 activation just overrides its relatively more efficient cell-level function which potentially makes an operative contribution to the functional dominance of CTLA-4 in the tumor microenvironment. Simulation results showed good agreement with data for tumor cells reduction and active immune cells count observed in each experiment.

These analyses propose a new mechanistic view on relative immunogenicity of PD-1 and CTLA-4 inhibitors manifested in literature and point the possible inherent obstacles in checkpoint inhibition-based immunotherapy of cancer to address in the future.

**Significance:** *Ex vivo* monitoring of temporal response to PD-1 and CTLA-4 in the closure of T cell movement dynamics and elucidating their feasible commitment to the kinetic constraints at cell-level resolution. Delayed dynamics of migratory response to CTLA-4 inhibition revealed a mechanistic view on potential T cell reinvigoration following immune checkpoint blockade.

## Introduction

Immune regulatory mechanisms modulate immune response, primarily to allow immune recovery and quench autoimmune reactions. Several mechanisms are involved in immune tolerance and malignant tumors use these mechanisms to escape immune rejection. Among them, immunosuppressive signaling pathways have a critical role in the regulation of chronic immune responses (1). Cytotoxic T-lymphocyte–associated antigen 4 (CTLA-4) and programmed death 1 (PD-1), two negative co-stimulatory receptors, attenuate T cells activation mainly through intrinsic cellular mechanisms (2).

The CTLA-4 molecules effectively blocks the CD28-depedent co-stimulatory pathway through competitive inhibition of ligands CD80 and CD86 binding thanks to its higher affinity for CD28. CD-28 dependent signals are mainly mediated by PI3K and Grb2 signaling molecules which have established contributions to actin-based cell movement alongside other effector functions (3–10). Besides, ligand-bound CTLA-4 by itself releases an inhibitory signal that interrupts CD28 and TCR molecular cascades (2, 5).

PD-1 engagement by its private ligands (PD-L1 and PD-L2), directly suppresses TCR signaling cascade through dephosphorylation and deactivation of its coupled components following the recruitment of specific protein-tyrosine phosphatases. However, most of the affected molecules are also involved in the regulation of cell migration, actin polymerization, and T cell anergy (10– 16). In addition, evidence suggests that PD-1 and CTLA-4 converge on the modulation of CD28 signaling pathway (17, 18).

Blockade of PD-1 and CTLA-4 receptors eliminates their downstream signals, which enables T cells reinvigoration and boosts antitumor response. The clinical success of checkpoint inhibition-based cancer treatment, is beholden to the fundamental research that provides a mechanistic understanding of the immune regulation and tolerance. However, improving therapeutic outcomes in Pd-1 and CTLA-4 blockade requires an even more detailed mechanistic insights particularly on comparative aspects of checkpoints function, which emerge from the coordinated nature of immunoregulatory mechanisms and the compensatory relationship between them (2, 19–21).

Despite the shared objective for PD-1 and CTLA-4 pathways, many studies indicate some peculiar specificities of each pathway; for example, in the timing of these receptors function, their corresponding immune cells population, the predominant operating environment, and the downstream transducing molecules (22–26). While TCR activation upregulates both receptors expression, the existence of CTLA-4 cytoplasmic reserves donate to this receptor the exclusive possibility of rapid intracellular trafficking to adjust its membrane recruitment according to TCR signal strength (27–29). These molecular and mechanistic characteristics can eventually appear in the dynamics of tumor-T cell interaction., However, in spite of the potential clinical relevance, comparative studies characterizing the kinetics of checkpoint induced immune inactivation, remained rare.

Several previous studies considered the dynamics of tumor-immune interaction in the presence of immunosuppressive pathways inhibitors for model-based investigation of their therapeutic outcomes(30–34). For example, a mathematical framework provided justifications for observations such as sustained tumor rejection in combined radiotherapy and checkpoint blockade despite tumor recurrence in exclusive regimens. The model highlighted their synergistic effect based on the immunogenicity of irradiation and intensification of abscopal effects in concurrent treatment (31). Also, an experiment-based mathematical model was used for effective dose prediction and optimal sequential strategy for radiotherapy and anti-PD(L)-1 treatment (32). While most of the models consider cell-cell interaction networks, the integration of related molecular processes can elucidate the relative effects of intracellular peculiarities and intercellular communication in response to immune checkpoint inhibition.

Herein, we aim to resolve the kinetic pattern govern PD-1 and CTLA-4 functions to address the possible therapeutic limitation originated from differential cell level response to these pathways.

For this purpose, we used a previously developed *ex vivo* system that recapitulate the tumor microenvironment and makes possible the precise monitoring of the tumor response to immune checkpoint blockade (35, 36). The system made up of a 3D microfluidic culture of organotypic tumor spheroids derived from patient samples that retain autologous immune cells (Fig.1a). The applied microfluidic device allowed for the controlled medium treatment with anti-PD-1 and anti-CTLA-4 antibodies as well as short-term evaluation of response to these inhibitors. Previously published data, based upon cytokine measurement and immune profiling of tumor spheroids, confirmed the above system as a novel platform for biomarker identification and systematic evaluation of checkpoint blockade outcome (37).

**Figure 1.**
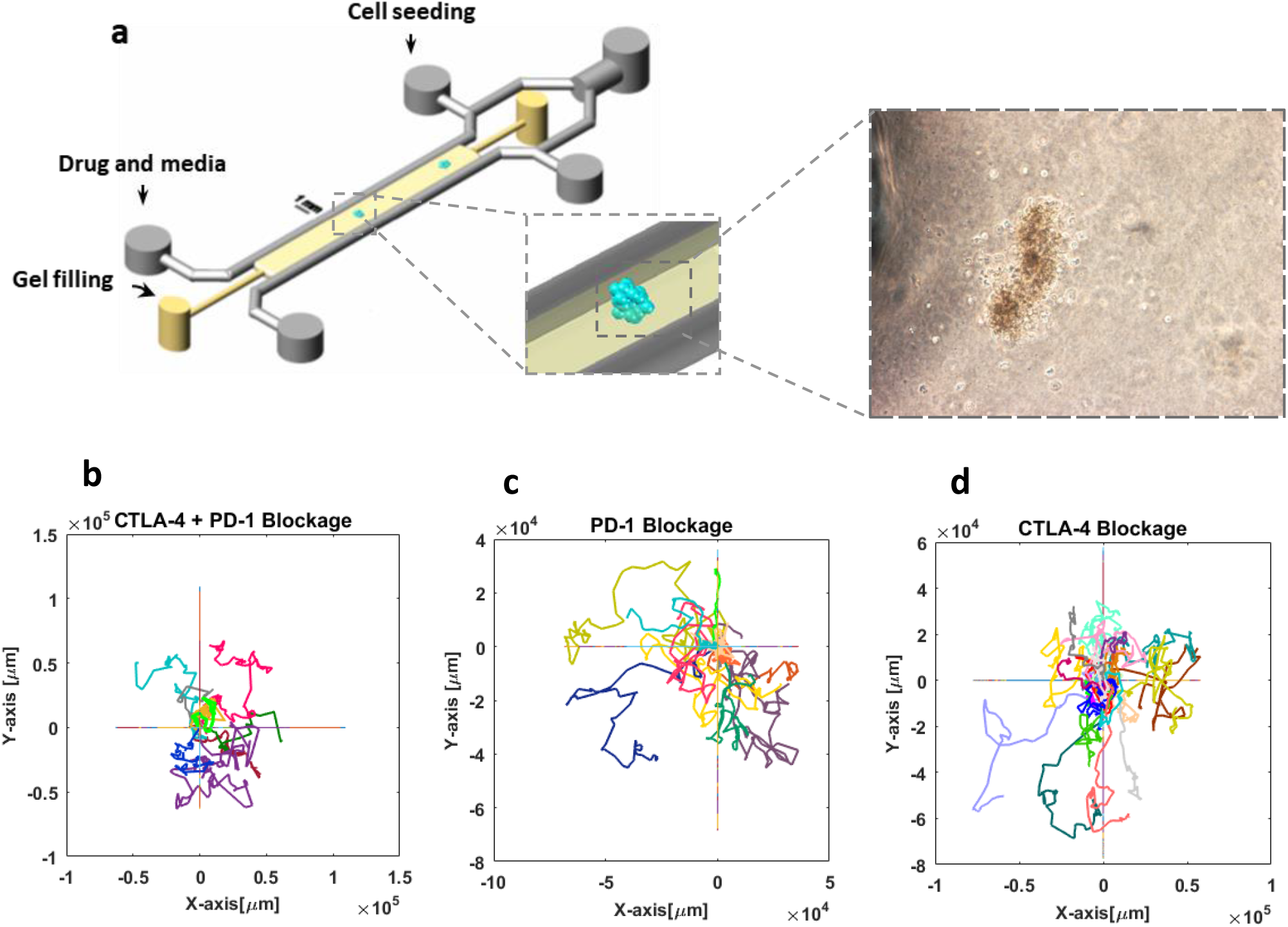
Tracking lymphocyte migration in 3D organotypic tumor culture. (a) Schematic view of the microfluidic device used for 3D culture of thyroid tumor spheroids (Scale bar shows 1 mm). Patient-derived tumor spheroids containing autologous immune cells are loaded in collagen medium in the central channel of the device and subjected to anti-PD-1 or anti-CTLA-4 antibodies treatment A representative bright-field image of a thyroid tumor spheroid shows the cells migrated into surrounding collagen. (b-d) Cell trajectories in three different conditions of the pathways inhibition. 12, 28, and 9 migrating cells were tracked in the condition of anti-PD-1, anti-CTLA-4 and combination exposure, respectively. For each experiment, Individual trajectories are shown with distinct color, and all trajectories started from the same point (coordinate center) that overlaid on the same graph. Time laps images captured every 15 minutes for at least 130 hours.

Here, using time-lapse imaging and single-cell tracking, time series of immune cell movements extracted in three conditions of individual and combined PD-1 and CTLA-4 blockade, to derive the state of immune cells activity and follow temporal response to immunosuppressive factors. The retrogressive movement capability of immune cells, as well as distinct dynamics of their response to PD-1 and CTLA-4 pathways, delicately captured by temporal parameter changes of a heterogeneous random walks model. Finally, a model of tumor-immune interaction based on a system of ordinary differential equations revealed how characteristic dynamics of PD-1 and CTLA-4 activation potentially imposes limitations on tumor response to PD-1 blockade.

## Methods

### Cell migration tracking

Patient derived organotypic tumor spheroids were studied for migration activity of immune cells in response to checkpoint blockade. Three and 10 spheroid cultures (biopsies form melanoma patients responsive to PD-1/PD-L-1 blockade) were investigated respectively for anti-CTLA-4 + anti-PD-1 treatments and exclusive inhibition of PD-1. Response to CTLA-4 was analyzed in five thyroid tumor cultures derived from a patient nonresponsive to PD-1/PD-L-1 blockade, the same sample was also accounted for six control experiment with no drug treatment. The detectable moving cells were tracked in each experiment (Supplementary Note 1) to extract cell trajectories, and related quantities such as cell velocity and persistence using the Tracking package of IMARIS software.

### Calculation of mean squared displacement of the cells

Mean squared displacement was evaluated as a substantial criterion to investigate the diffusive behavior of the cells. The MSD of a walker is defined as follows:

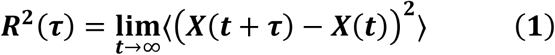

Where |***X*(*t* + *τ*) − *X*(*t*)**| corresponds to the displacement of the walker between two consecutive steps. The parameters ***τ*** and ***t*** refer respectively to the time interval and total time of the movement. The MSDs were calculated for the cell trajectories as well as simulated ones to check their correspondence and evaluate the model of migration.

### Bayesian inference method for parameter estimation of the migration model

We used heterogeneous random walks(38) for modeling the migration of tumor-infiltrating (Antigen-experienced) lymphocytes and detection of temporal variations of model parameters affected by immune checkpoint activation. We applied an algorithm designed and implemented by Metzner, C. et al.(38) for sequential estimation of the random walks model parameters, based on Bayesian inference method. Accordingly, the displacements are calculated using cell velocities in each time step, and the time-varying displacement vector is described according to a first-order autoregressive model:

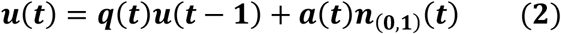

The parameter ***q*(*t*)** represents the time-varying persistence of the cells ranged between −1 to 1 from anti-persistent to persistent random motion. The parameter ***a*(*t*)** corresponds to the motion activity, namely the intensity of noise in the random walk process. The random noise vector ***n* = (*n***_***x***_ **; *n***_***y***_) is taken from an uncorrelated Gaussian distribution with unit variance. The likelihood function of the stochastic process conducts the sequential Bayesian method to infer the joint distribution of the parameters for each time step. Single cell activity and persistence are given in Supplementary Fig. S2.

### Simulation of cell trajectories

The time-varying autoregressive model of the first order was evaluated for analysis of cell movements. In this way, we used the estimated activity and persistence of the cells to simulate trajectories based on the same autoregressive process. The model parameters were further estimated for the simulated trajectories to be compared with the input parameters and confirm the accuracy of the applied algorithm. The later statistical verification of the simulated trajectories determines the accuracy of this modeling paradigm.

### Mathematical Modeling

A set of biological assumptions have been considered to infer a mathematical model of checkpoint induced immune modulation. These assumptions outline the theoretical underpinning of the work and reduce the parameters and equations to those sufficient for rational description of the experimental data.

### Model hypothesis

#### Accessibility of tumor cells

The experimental results indicate that even in the case of two pathways blockade a fraction of tumor cells would survive. We supposed that this is because of the limited access of tumor infiltrating lymphocyte to the population of tumor cells, a well-known phenomenon play a role in tumor-associated immune resistance.(39) Assuming a spherical geometry for tumor spheroids, equation (7) relates the accessible fraction of tumor cells to their total population(40).

#### Temporal delay in surface expression of PD-1 receptor

Most of the cellular CTLA-4 receptors are embedded within cytoplasmic vesicles. Surface expression of CTLA-4 is quickly upregulated by membrane trafficking control of this intracellular pool. In contrast, genetic and epigenetic mechanisms are mainly engaged in the regulation of PD-1 expression which leads to a time delay in T cell inactivation induced by the PD-1 pathway (27–29).

#### Direct and indirect tumor - T-cells interaction

The subpopulation of accessible tumor cells may be directly killed by T-lymphocytes and in turn, stimulate PD-1 and CTLA-4 pathways via direct contact. The effects of T cell exhausting factors other than PD-1 and CTLA-4 (i.e. free radicals, secreted cytokines, oxygen limitation, …) are assumed to be dependent on the total population of tumor cells (regardless of the direct accessibility to T cells) (41, 42).

#### Up-regulation of PD-1 expression

We assume that sustained presence of cancer cell antigens in the cultured tumor microenvironment leads to the up-regulation of PD-1 surface expression. The rate of this induction is considered to be independent of cancer cell populations.

#### No proliferation and recruitment in 3D channel microenvironment

No immune cell recruitment occurs in this experimental setup, and tumor cell proliferation can be ignored because of limited nutrient availability in this condition.

#### Lymphocytes activity modeling

It is assumed that tumor cell death occurs in proportion to lymphocytes activity which appears in their migratory behavior. Therefore, the dynamics of activity in tumor interacting lymphocytes is modeled, instead of their population changes.

### Dynamical modeling of tumor infiltrating lymphocytes inactivation induced by CTLA-4 and PD-1 signaling pathways

As stated above genetic, transcriptional, and translational regulation of PD-1 expression cause a few hours’ delay in lymphocyte inactivation triggered by this receptor. Considering the above assumption, along with other model hypothesizes, and taking into account PD-1, CTLA-4, and other immunosuppressive factors in the tumor microenvironment, the dynamical model of tumor cells-T cells interaction is schematically represented in Fig. 2.

**Figure 2.**
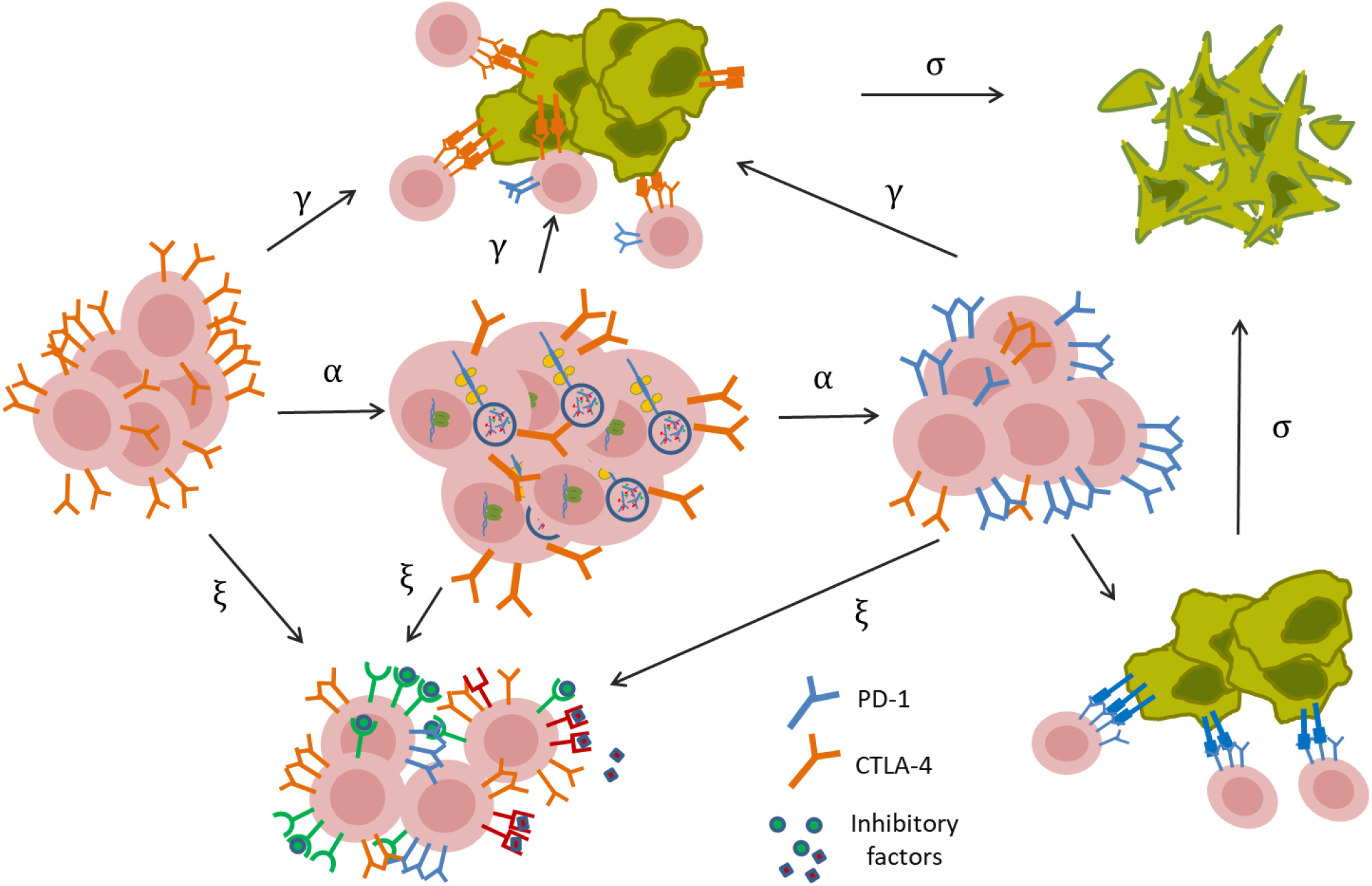
Illustration of the relationships between the components of the mathematical model. PD-1 and CTLA-4 receptors and other factors suppress lymphocytes activity in the tumor microenvironment. As a result of the cytoplasmic storage of preformed CTLA-4 molecules, the total population of active lymphocytes can express CTLA-4 receptor with no time delay and can be potentially inactivated by this receptor with gamma rate. Other inhibitory factors as well, can suppress lymphocytes, independent of PD-1 receptor. Regulation of PD-1 expression mainly at genetic and epigenetic levels, results in a time delay in immune suppression triggered by PD-1 receptor. Concurrently, the cytotoxic activity of lymphocytes resulted in the death of tumor cells.

According to PD-1 expression state, the dynamical model comprises activities for three subpopulations of lymphocytes: (1) lymphocytes with unexpressed **(*L***_***u***_), (2) partially expressed **(*L***_***p***_), and (3) fully expressed **(*L***_***e***_) PD-1 receptors. The subpopulation of lymphocytes lacking PD-1 receptors converts to PD-1 expressing cells with a delay of second-order and expression rate ***α***. The conversion then is followed by PD-1 induced inactivation with ***β*** rate in contact with the accessible tumor cells (Eq. 3-6).

It is assumed that all of the three subpopulations of active lymphocytes may be deactivated by CTLA-4 receptors with *γ*rate along with inhibitory factors other than PD-1 and CTLA-4 (i.e., radical formation, hypoxia …) with **ξ** rate of performance. PD-1 signaling pathway can exclusively inhibit the cells in full expression state of PD-1 **(*L***_***e***_). Cytotoxic activity of lymphocytes total population causes the death of tumor cells at a rate of σ (Eq. 6).

Finally, the presumption of cell to cell contact in CTLA-4 and PD-1 pathways activation as well as tumor cell death process leads to a system of ordinary differential equations as follows:

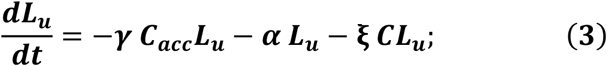

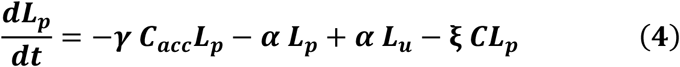

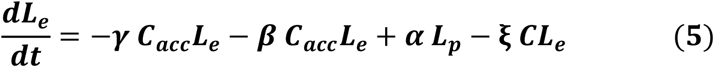

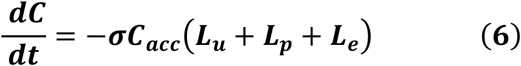

Where the accessible part of tumor cells (***C***_***acc***_) is calculated from the total population (***C***) as follows:

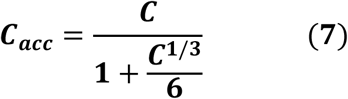

This equation is derived by assuming a simple spherical geometry for the tumor spheroids and subsequent calculation of the cells which hold surface volume (40).

## Results

### Stochastic modeling of T cells migration

#### T cells trajectories

The instances of T cell trajectories under the condition of PD-1 and CTLA-4 blockade alone or in combination are illustrated in Fig.1. The tracks represent 2D projection of cell trajectories in 3D cell cultures. These results show temporal variations of T cell step length and for most of the cells, these variations display a decreasing trend in the time series of cell movements.

#### Mean squared displacement of the cells

It was observed that the slope of the MSD_log−log_ plots of the cells takes values smaller than one at many time intervals, which is a characteristic of sub-diffusive motions. Furthermore, in all of the three experimental conditions, the slope of the curves has a decreasing trend and almost tends to zero (Fig. 3d-f, blue curves).

**Figure 3.**
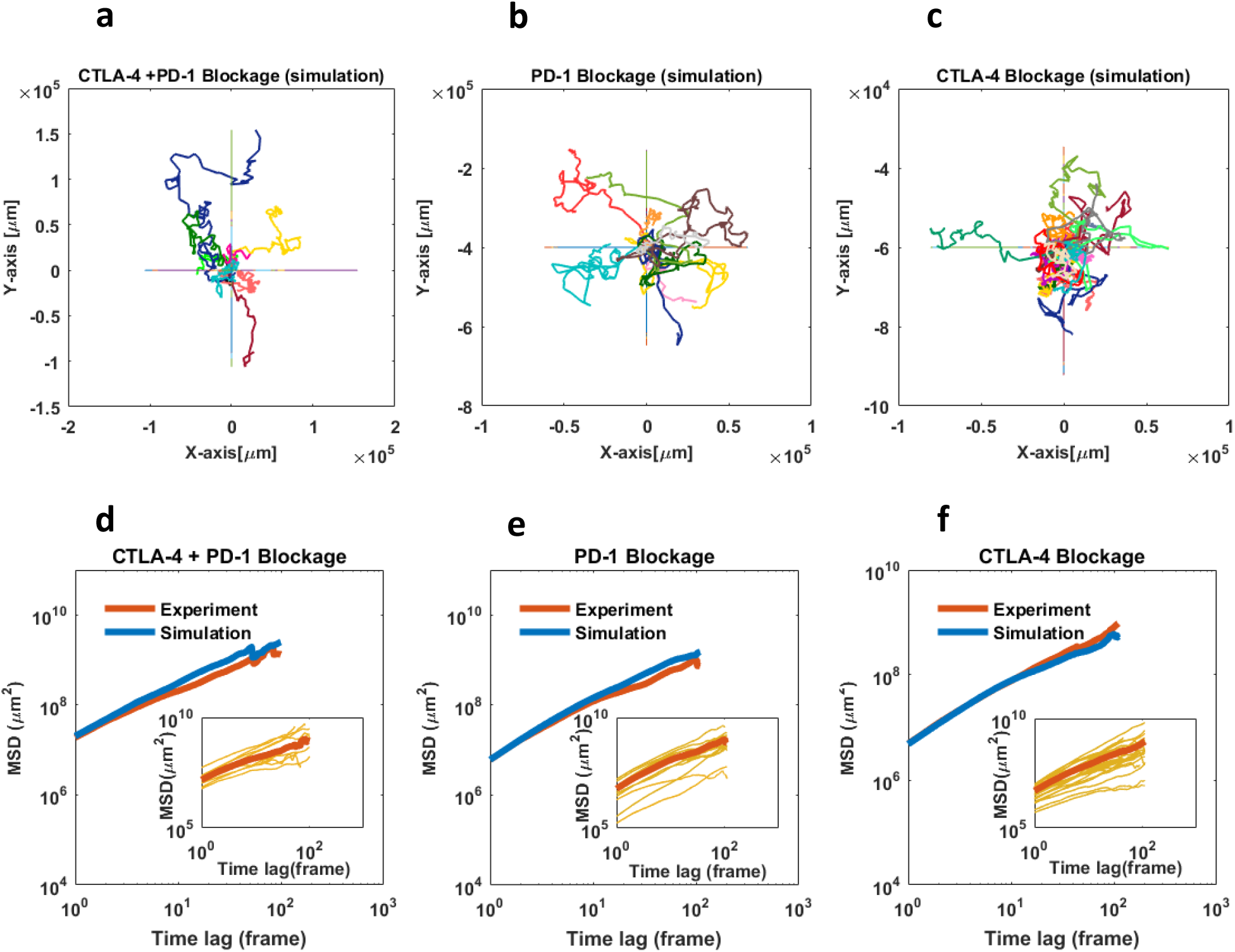
Statistical analysis of cell migration and AR-1 model evaluation. (a-c) Simulated trajectories using AR-1 model with time-dependent parameters (activity and persistence) based on single-cell parameters change estimated from experimental data. (d-f) Mean squared displacements (population average) of tumor infiltrating lymphocytes treated with anti-PD-1 or anti-CTLA-4 antibodies, or a combination thereof (blue curves). The MSDs show sub-diffusive behavior with the approximate power-law exponent of 0.83 for PD-1 0.93 for CTLA-4 and 0.98 for combined inhibition. Red curves represent the MSDs of simulated trajectories for each experiment. The intercepts of the curves indicate the highest value of cell diffusion coefficient in combined treatment (D = 7.3). AR-1 model with changing parameters reproduce well the experimental MSDs obtained from each condition. Insets: MSD curves for individual cells. The last steps of the migration trajectories removed because of error amplification in MSD calculation.

#### Temporal variation of cell migration parameters in checkpoint blockade experiments

We supposed that the observed migratory behavior can be explained by a model of random walks with time-dependent parameters. Through this assumption, we used a *first*-*order autoregressive model* with time-varying parameters to model the studied cells movement. Sequential Bayesian inference method was applied on the cell trajectories to deduce the statistical parameters over the single-cells migration period (Supplementary Note 2).

For evaluation of the autoregressive (AR-1) model of migration, the estimated parameters (time-varying persistence and activity of the cells) were used in an inverse manner to see if the model reproduces trajectories that statistically match the migration data (Fig. 3a-c). In this way, the Bayesian method and applied algorithm were primarily assessed for simulation of trajectories able to conserve the input parameters (Supplementary Fig. S2). Hereon, the MSDs of the simulated trajectories would show if the AR-1 model with changing parameters captures the migratory behavior of the cells. As shown in Fig. 3d-f, there is a good agreement between population-averaged MSDs of the simulated time series and experiment-derived ones.

Temporal pattern of movement activity and persistence of the cells are illustrated in Fig. 4a-c. In each experiment, while the activity parameter shows considerable changing behavior, population-averaged persistence parameter displays moderate variations over the time of observation (Fig. 4a-c, insets). In the case of PD-1 blockade, ensemble activity of the cells displays a relatively constant decrease over time from the very beginning, while blocking the CTLA-4 receptor, drives the cells to a more rapid decrease of the motion but after a short initial delay (Fig. 4b, c). In the case of drug combination, the migratory activity of the cells showed more intensive fluctuations but only a mild overall decrease during the time course of the experiment (Fig. 4a). Additionally, as observed from time-zero values of the activities, the cells more intensely retrieved their motion strength following CTLA-4 blockade (compared with PD-1 blockade). This initial activity did not decrease significantly in combined receptor blockade while in both cases of exclusive receptor inhibition, activity reduction continues to approximately reaches zero value.

**Figure 4.**
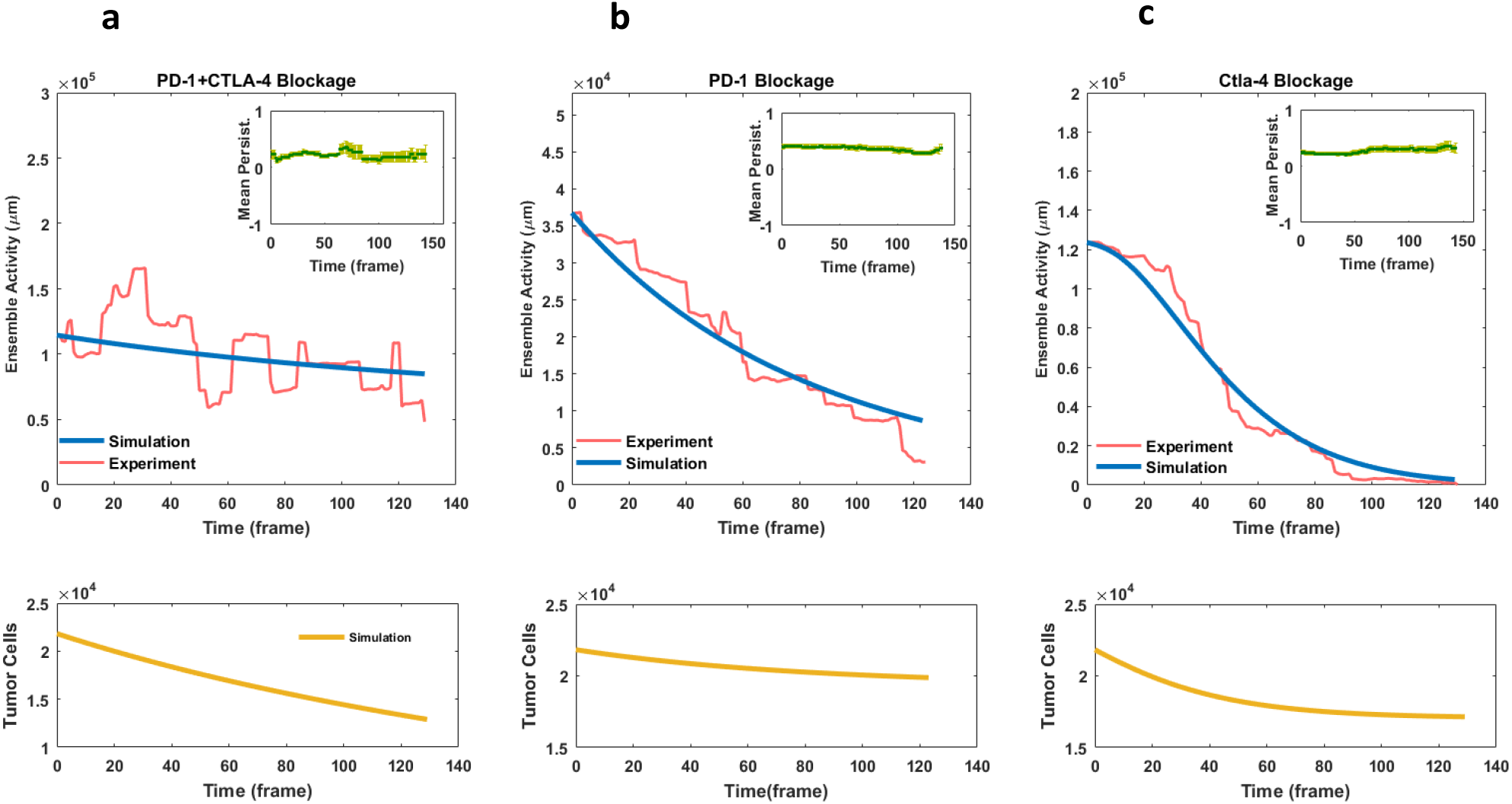
Temporal variation of immune cells activity and dynamical model simulation in (a) combined inhibition as well as exclusive blockade of (b) PD-1 and (c) CTLA-4 pathways. Immune cells activity displays a much lower initial value in PD-1 blockage compared with two other experiments. Contrary to PD-1 blockage the decrease in cellular activity in CTLA-4 blockade entails a delay of several hours, while in the combined inhibition of the pathways, the cells remain active up to the last frames of the assays. Dynamical model parameters were inferred by fitting the model (blue curves) to immune activity change derived from cell migration data (pink curves). Tumor-immune profiling of the spheroids as well as live-dead cell staining confer the initial value of 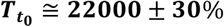 and final values of 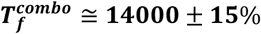 and 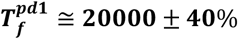 and 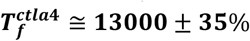 for tumor cell population (yellow curves). Simulation results with specified parameter set (σ = 0.00000022, ξ = 0.000000137, γ = 0.0000024, α = 0.0549 and β = 0.000016; all in a temporal unite of time-step) show good agreement with relative tumor killing performance of anti-PD-1 and anti-CTLA-4 components according to cell death assessments in tumor spheroids. No unit considered for cellular activity. Inset: Population-averaged persistency of immune cells migration in checkpoint inhibition experiments. The average values of persistence parameter show moderate changes over the time of experiments. Error bar corresponds to standard error.

### Dynamical modeling of lymphocytes activity in the presence of immune checkpoints inhibitors

Dynamics of the inferred statistical parameters was investigated to elucidate cellular response to PD-1 and CTLA-4 checkpoint inhibitors reflected in lymphocytes moving behavior. Accordingly, immobilization of immune cells in the tumor spheroids was leveraged to characterize tumor-induced T cell suppression in tumor microenvironment (see Method section). No significant cell movement was observed in cultures without drug treatment (control samples, see supplementary information). The experimental data for the number of alive tumor cells was used to evaluate the proposed model.

#### Model simulation for the combination of PD-1 and CTLA-4 blockade

As schematically illustrated in Fig. 2, in the absence of PD-1 and CTLA-4 signaling cascades, other inhibitory signals and factors in tumor microenvironment modulate lymphocytes activity with decreasing rate of *ξ*. In addition, the cytotoxic activity of T cells decreases the population of tumor cells at a rate of σ.

It was assumed that the whole population of experimentally measured spheroid tumor cells (≅ 22^′^000 cells ± 30%, Supplementary Note 1) are initially alive. Furthermore, the initial value for T-cells activity corresponds to the relevant value obtained from the autoregressive model at time 0. To infer the dynamical model parameters, we applied quasi-Newtonian algorithm using unconstrained Nonlinear programming solver of MATLAB optimization toolbox (MATLAB R2016a, MathWorks). Setting α, β, and λ parameters to zero (Eq. 3-6, Method section), the parameters ξ and σ were determined, so that simulation results reproduce the final tumor cell count as well as immune activity change in this experiment (Fig. 4a, blue and yellow curves). The optimal solution was found at the value of 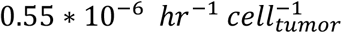 for lymphocytes inactivation rate (ξ), caused by factors other than the two blocked receptors. Moreover, considering the experimental value for the final tumor cells population ≅14’000 cells ± 15%, tumor cell death rate (σ) was estimated to be 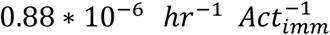.

#### Model simulation for PD-1 blockade

In the absence of CTLA-4 inhibitor, activation of the relevant signaling pathway affects the dynamics of lymphocytes activity (Fig. 2).

In this scenario, the parameters ξ and σ set as the values inferred from the previous simulation and similarly the initial values experimentally determined for this condition. In addition, tumor cell death data are used for model validation.

Training the experimental diagram, the value of 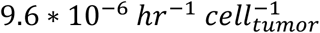 was estimated by the solver for lymphocytes inactivation rate triggered by CTLA-4 signaling cascade (***γ*)** (Fig. 4b, blue curve). As a model validation, the dynamical model in this parameter adjustment would capture tumor cell death data obtained from PD-1 blockade experiments (compare the experimental value of 20^′^000 cells ± 40%; with model outcome of 19’500 cells for the final tumor cells population, Fig. 4b, yellow curve).

#### Model simulation for CTLA-4 blockade

In this case, up-regulation and consequent activation of PD-1 pathway modifies the dynamics of lymphocytes activity (Fig. 2). To set the initial values of the model we supposed that the whole population of activated lymphocytes initially belongs to the subpopulation lacking PD-1 receptor. In this standing, the parameters ***α*** and ***β*** were learned so that the model could explain the experimental data (Fig. 4c, blue curve).

Because of the possibly unrealistic assumption above, we attempted to correct the initial condition of the model to get a more accurate estimation of ***α*** and ***β*** parameters value. Details are given in Supplementary Note 3.

Simulation results showed that the model is capable of reproducing inactivation pattern of T cells for α = 0.22*hr*^−1^ and 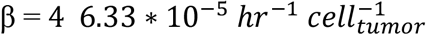 which is more than six fold greater than CTLA-4’s rate of function. The subsequent correction of baseline situation, made no significant improvement in the parameter values (Supplementary Note 3). The model was further validated by the relative concordance of the simulated cancer cells decline with tumor killing performance of CTLA-4 blockade (compare the final population of tumor cells in anti-PD-1 and anti-CTLA-4 treatments, Fig. 4c yellow curve). The inferred values of the parameters represented in Table 1.

**Table 1.** Tumor immune interaction parameters.

| *Parameter* | *Definition* | *Optimal Value\** | *Unit\*\** |
| --- | --- | --- | --- |
| $\sigma$ | Tumor cell death rate | $8.8 * 10^{-7}$ | $hr^{-1} Act_{imm}^{-1}$ |
| $\gamma$ | CTLA-4 induced immune cells inactivation rate | $9.6 * 10^{-6}$ | $hr^{-1} cell_{tumor}^{-1}$ |
| $\beta$ | PD-1 induced immune cells inactivation rate | $6.33 * 10^{-5}$ | $hr^{-1} cell_{tumor}^{-1}$ |
| $\xi$ | Non-checkpoint induced immune cells inactivation rate | $5.5 * 10^{-7}$ | $hr^{-1} cell_{tumor}^{-1}$ |
| $\alpha$ | PD-1 expression rate | 0.22 | $hr^{-1}$ |

\* Optimal values estimated by fitting model simulation to corresponding experimental results.
\*\* Accounting for the frame time interval corresponds to 15 min and unit-free measure for cell activity.

#### Comparing in vivo population of reinvigorated cells in PD-1 and CTLA-4 blockade

In addition to the investigated spheroids (average size of 70 µm (37)) we ran the model for a more realistic tumor size about 100 *mm*^3^ (37) better mimicking in vivo condition and the consequent therapeutic implications of the model.

Results demonstrate that the population of immune cells deactivated by CTLA-4 receptor, by nearly ten-fold, out-numbers the cells whose inactivation associated with PD-1 receptor. Presuming that this model outcome is a good representation of the evolved tumor situation, this suggests that the inhibition of the CTLA-4 pathway would reactivate a larger population of immune cells than PD-1 pathway blockade.

Table 2. represents the average count of the cells which are set to motion following each treatment scenario. These data indicate that CTLA-4 blockade in patient-derived tumor spheroids is capable of reinvigorating a larger population of lymphocytes, that is in good agreement with the model prediction. Indeed, because of defined time delay in PD-1 suppressive function, sufficiently large population of tumor cells drive immune cells to the inactivated state primarily and more efficiently by CTLA-4 receptor stimulation and signaling.

**Table 2.** Reinvigorated cells count in checkpoint blockade experiments.

| *Type of Treatment* | *Spheroids Active Cells Number (Average)* |
| --- | --- |
| **Anti- CTLA-4** | **$65.6 \pm 9.38$** |
| **Anti-PD-1** | **$19.3 \pm 1.64$** |
| **Combo** | **$61.6 \pm 7.75$** |

## Discussion

T cell migration can be modulated by PD-1 and CTLA-4 signaling pathways through molecular intermediates which regulate actin polymerization, cytoskeletal structure and membrane organization (6–10, 13–16), as supported by the observation that immune cells displayed increased and sustained motility in the presence of combined pathway inhibitors. Although, such a stimulatory effect on T cell trafficking converges with the other invigorating cellular events following checkpoints inhibition, the sub diffusive behavior of the cells even at the beginning of the assessment, demonstrates T cells tendency for physical engagement rather than pointless receding movements in tumor environment, (Fig. 3,4).

In the current study, the motion pattern of tumor-associated immune cells was investigated in 3D *ex vivo* culture, and their motion behavior leveraged for the dynamical modeling of tumor-immune cell interaction dominated by immunosuppressive pathways. To this end, time series of cell migration events were applied to infer the extent of immune modulation related to each pathway; using this functional response, immune cells inactivation was quantified and the relative kinetics of immune suppression induced by PD-1 and CTLA-4 receptors was modelled.

As shown in the activity diagrams, T cell movement affected by PD-1 and CTLA-4 pathways depicts distinct profiles of variation such that T cells mobility behavior dominated by PD-1 pathway (anti-CTLA-4 treatment) represents a significant time lag before a continuous decline, which we presumed to be a time imposition for de novo receptor expression.

The proposed dynamical model was able to explain distinct immunostimulatory effects of PD-1 and CTLA-4 pathways based on different mechanisms involved in the regulation of their expression and function.

Experiment-based estimation of the parameters revealed the greater potency of immune modulation for PD-1 rather than CTLA-4 receptor (see Table 1), in agreement with the previous findings demonstrated that PI3K/Akt signaling transversed by PD-1 and CTLA-4 pathways employing different molecular networks which is more effective in PD-1 signaling cascade (25).

Importantly, the model could explain how, despite the functionally more efficient signaling cascade initiated by PD-1, the presence of cytoplasmic vesicles store CTLA-4 receptor, eventually leads to a more dominant immunosuppressive effect of this receptor at the level of lymphocyte populations (43).

The model assertion that the specific regulation of CTLA-4 expression, mainly relying on vesicular transport, may precipitate its suppressive function, has been acknowledged by a study aimed at the modeling of CTLA-4 intracellular trafficking, showing the significance of CTLA-4 recycling in the kinetics of its molecular function. They show that even in the presence of high-affinity ligands, the intracellular CTLA-4 resources, lasts for several hours thanks to its cytoplasmic reserves and only thereafter its surface expression turns to be dependent on de novo synthesis (44).

Furthermore, the key assumption linking the more limited function of PD-1 to its delayed expression as a consequence of transcriptional regulation, is well supported by the studies show evidences for a profound regulation of PD-1 expression and divide lymphocytes population to the subpopulations displaying distinct level of PD-1 expression (45, 46).

Our analyses suggest that the relative population inhibited by these two pathways, depends on the population of tumor cells. When the tumor size is relatively large, time delay in PD-1 surface expression leads to the broader inactivation of immune cells due to CTLA-4 function. and potentially shapes an inherent limitation for anti-PD-1 treatment outcome. Consistent with this notion, baseline tumor size may be considered as a candidate biomarker of response to anti-PD-1 therapy (47, 48). While the antigen burden (i.e. tumor mutation burden) is already considered as a possible explanation for tumor-size effect, our study suggests that even in spite of pretreatment adaptation of tumor-immune populations, the marker of tumor size, remains as a limiting factor in patient response to anti-PD-1 treatment (49, 50). These results are in consistence with the studies show a broader immune restoration and more significant TCR repository enrichment in mature tumors following CTLA-4 inhibition (51).

It should be noted that this relatively broad immunogenicity may lead to the activation of other immunosuppressive pathways and a more intense reinvigoration of regulatory cells, possibly resulting in the limited therapeutic response to anti-CTLA-4 compared with anti-PD-1 treatment (43, 51–56). Indeed, our analyses further remark the importance of pretreatment immune profiling and highlights the significance of identifying predictive biomarkers as a strict irrevocable goal for improving response to CTLA-4 blockade(43, 55).

On the other hands, considering the importance of the initial response and priming of anti-tumor immunity for immunotherapy outcome, this profound immune excitation predicted here provide some rationale for combined and alternative therapeutic strategies; patient treatment with immunostimulatory drugs with lower toxicity following initial activation of the immune system by CTLA-4 blockade may improve clinical outcomes(57). Moreover, in patients without baseline population-related biomarkers for anti-PD-1 treatment, this strong immune restoration may establish the condition for improved objective response (51, 54, 58).

The presented quantitative insight into the distinct PD-1 and CTLA-4 kinetics and functions can be useful for the development of more practical models for planning effective treatment schemes.

More generally this study highlights the need for more accurate understanding of the kinetics of response to checkpoints inhibitors in order to develop more effective therapeutic strategies based on sequential stimulation of the immune system.

## Notes

https://github.com/safaeifard/F-Safaeifard/tree/main/Heterogenous%20AR-1%20Model

### Competing Interest Statement

The authors have declared no competing interest.

