## Supplementary Information for "Delayed dynamics of migratory response to CTLA-4 blockade reveals a mechanistic view on potential T cells reinvigoration following immune checkpoint blockade"

### Supplementary Note 1

#### Experimental background

During the experimental procedure patient-derived tumor spheroids in combination with collagen were loaded into the central channel of 3D microfluidic culture device (Fig.1 main text). The cell culture was treated with monoclonal antibodies or their combinations. Pembrolizumab and Ipilimumab antibodies were used as PD-1 and CTLA-4 pathways inhibitors, respectively. The injected dose (clinically used 1:100 dilutions of stock concentrations) resulted in peak plasma concentrations measured following administration of 10mg/kg of each drug (FDA CDER application). In these experiments, tumor spheroids were targeted by different immune markers, and their immune profile was characterized via flow cytometry technique.
The presence of different cytokines and their level of concentration in the cell culture microenvironment were evaluated using a bead-based immunoassay approach. These outcomes of assays show the success of the designed cell culture to recapitulate tumor microenvironment considering its autologous cell population and chemical signals.
In addition, live/dead cell quantification was performed using dual labeling deconvolution fluorescence microscopy. The presence of lateral channel along the central area of the culture medium entails the possibility of controlled drug injection. Time-lapse imaging of cell mobility was carried out through the transparent cover of the chamber. All the experiments were performed at Dana–Farber Cancer Institute. You find detailed information in the article previously published by Jenkins, et al. *Cancer Discovery* 8.2 (2018).
In the following the experimental results, on the topic here are discussed.

##### Tumor samples

The immune cells were investigated in organotypic culture of tumor spheroid (including lymphoid and myeloid cells) derived from patient samples of thyroid and melanoma cancers. Depending on patient's response to ICB components, the spheroids were cultured in four types of medium containing: (1) PD-1 and (2) CTLA-4 inhibitors, (3) their combination alongside (4) control cultures.

##### Live-Dead imaging of tumor spheroids

With the goal of tumor and non-tumor cell death quantification, live/dead imaging was performed using dual labeling deconvolution fluorescence microscopy (Fig. S1), and results are shown in Table S1.

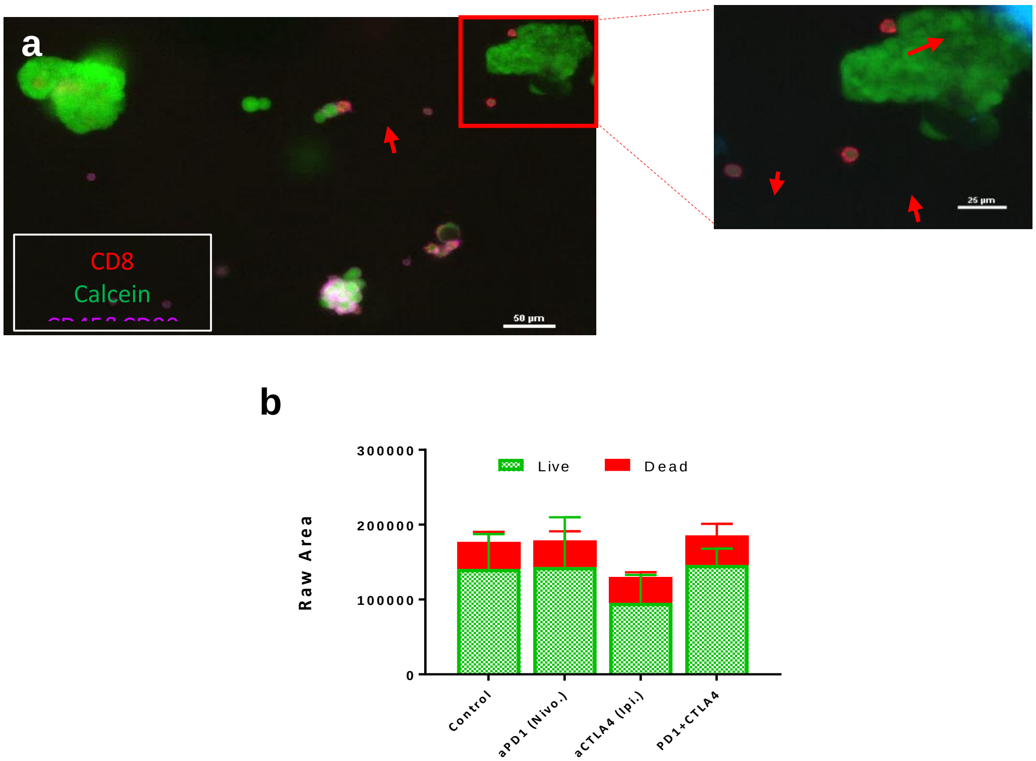


**Figure S1. Immune profiling and viability evaluation of patient-derived thyroid tumor spheroids.** (a) A representative image of immunofluorescence staining that identifies the population of autologous CD45+ and CD8+ immune cells in organotypic tumor spheroids derived from patient samples. (b) Live (AO = green)/dead (PI = red) quantification of tumor spheroids following culture treatment with Nivolumab (anti-PD-1), Ipilimumab (anti-CTLAA-4) and combination.

**Table S1. Spheroids live-dead and tumor-immune profiling**

| Experiment | Spheroids Population | **Tumor Cells Ratio*** | Tumor Death Ratio** |
|---|---|---|---|
| Control | **175000 ± 25%** | **0.22** | 0.090 |
| PD-1 | **175000 ± 30%** | **0.16** | 0.196 |
| CTLA4 | **130000 ± 25%** | **0.16** | 0.483 |
| Combo | **180000 ± 30%** | 0.13 | 0.433 |

*The ratio of tumor cells to total population of spheroid cells (Tumor + Immune cells) for each drug treatment
** The ratio of tumor cells to total tumor cells population of spheroids (Dead + Live cells)

These data were used to calculate the initial and final population of tumor cells ($TDR$: $Tumor\ Death\ Ratio$).

$$Tumor\ Population_{t=0} = Average\ Spheroids\ Popultion * Average\ Spheroids\ Tumor\ Cells\ Ratio * (1 - TDR_{control})$$

$$Tumor\ Population_{final} = Tumor\ Population_{t=0} - \frac{TDR_{treatment} - TDR_{control}}{1 - TDR_{control}} * Tumor\ Population_{t=0}$$

Which gives the initial and final values of tumor cells population:

$$T_{t_0} \cong 22000 \pm 30\%$$

$$T_f^{combo} \cong 14000 \pm 15\%$$

$$T_f^{pd1} \cong 20000 \pm 40\%$$

$$T_f^{ctla4} \cong 13000 \pm 35\%$$

**Time-lapse imaging of tumor culture**

Time-lapse images of bright field microscopy obtained during the experiments. The images were captured, every 15 minutes, using a Nikon Ti inverted microscope with a 10x NA 0.3 objective and cooled CCD camera (Orca R2, Hamamatsu) in a humidified, temperature-controlled chamber. Illumination was with a CoolLED pe-100 white light LED. Imaging of various fields of view overtime was controlled by NIS-Elements software of a Prior motorized stage along with the LED and camera. Image capturing was carried out for more than 30 hours (Fig.1, main text).

**Activated cell count in each treatment**

The number of lymphocytes that set to motion, was estimated at the beginning of PD-1 and CTLA-4 pathways blockage. For this aim, we used time-lapse microscopic images in the first few frames to count moving cells in microfluidic cell cultures treated by each blocking agent.

#### Supplementary Note 2

##### Evaluation of parameter estimation for AR-1 model of T cell migration

A sequential Bayesian inference method corresponding to autoregressive model of the first order, was applied on the cell trajectories to deduce the joint probability distribution of the model parameters for each time step (Fig.1 main text). We used the average values of the inferred distributions to trace statistical parameters change over the single-cell migration period.

To evaluate the method and the parameter estimation algorithm for our migration assay, single-cell time-varying parameters were used in an inverse manner to see if the model can reproduce trajectories that statistically match the migration data.

The results of parameter estimation for simulated trajectories indicated that Bayesian method and the applied algorithm are able to conserve the input temporal change of the model parameters in simulated trajectories (Fig. S2).

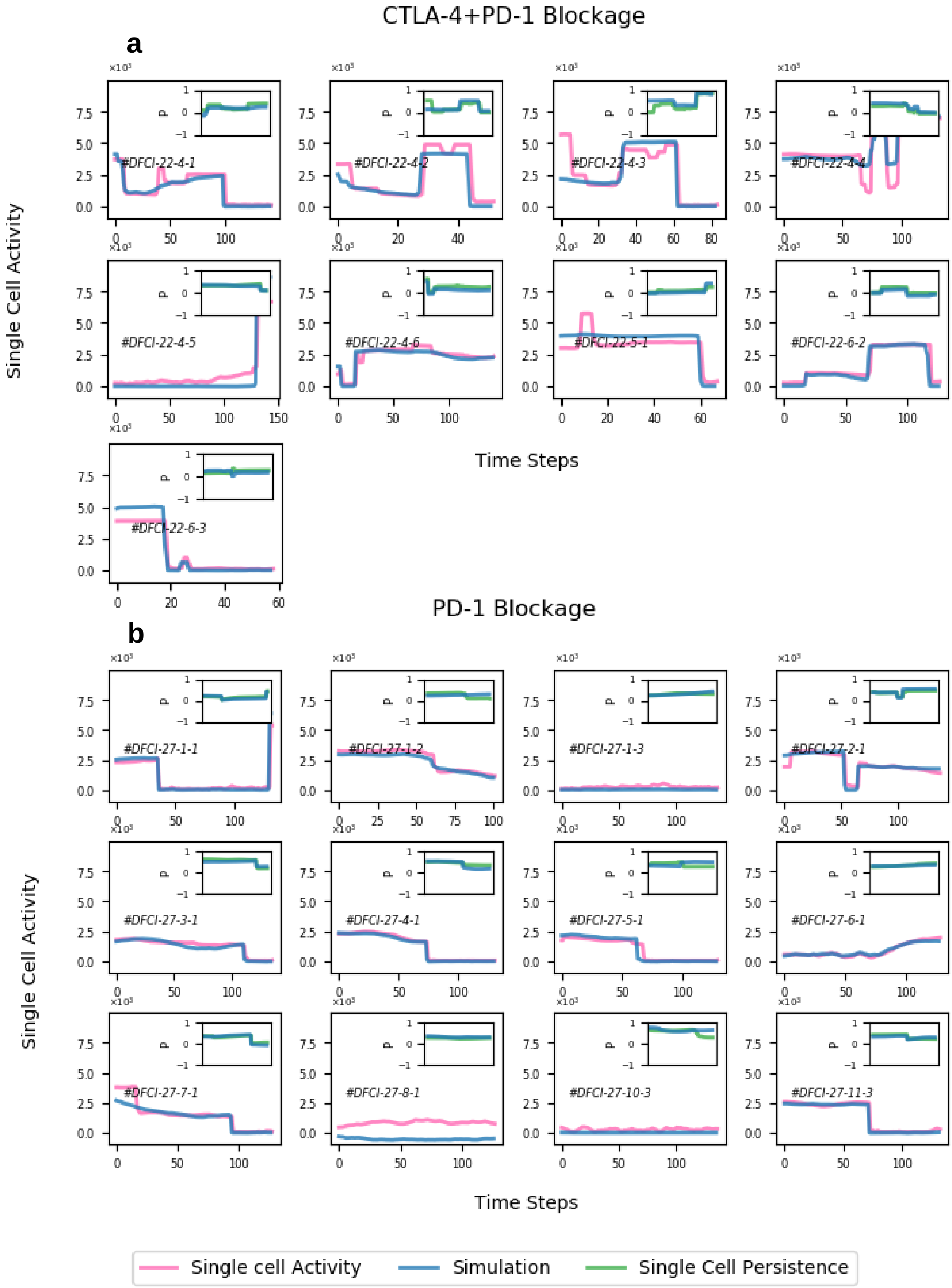
CTLA-4+PD-1 Blockage
a
#DFCI-22-4-1
#DFCI-22-4-2
#DFCI-22-4-3
#DFCI-22-4-4
#DFCI-22-4-5
#DFCI-22-4-6
#DFCI-22-5-1
#DFCI-22-6-2
#DFCI-22-6-3
Single Cell Activity
Time Steps
PD-1 Blockage
b
#DFCI-27-1-1
#DFCI-27-1-2
#DFCI-27-1-3
#DFCI-27-2-1
#DFCI-27-3-1
#DFCI-27-4-1
#DFCI-27-5-1
#DFCI-27-6-1
#DFCI-27-7-1
#DFCI-27-8-1
#DFCI-27-10-3
#DFCI-27-11-3
Single Cell Activity
Time Steps
Single cell Activity
Simulation
Single Cell Persistence

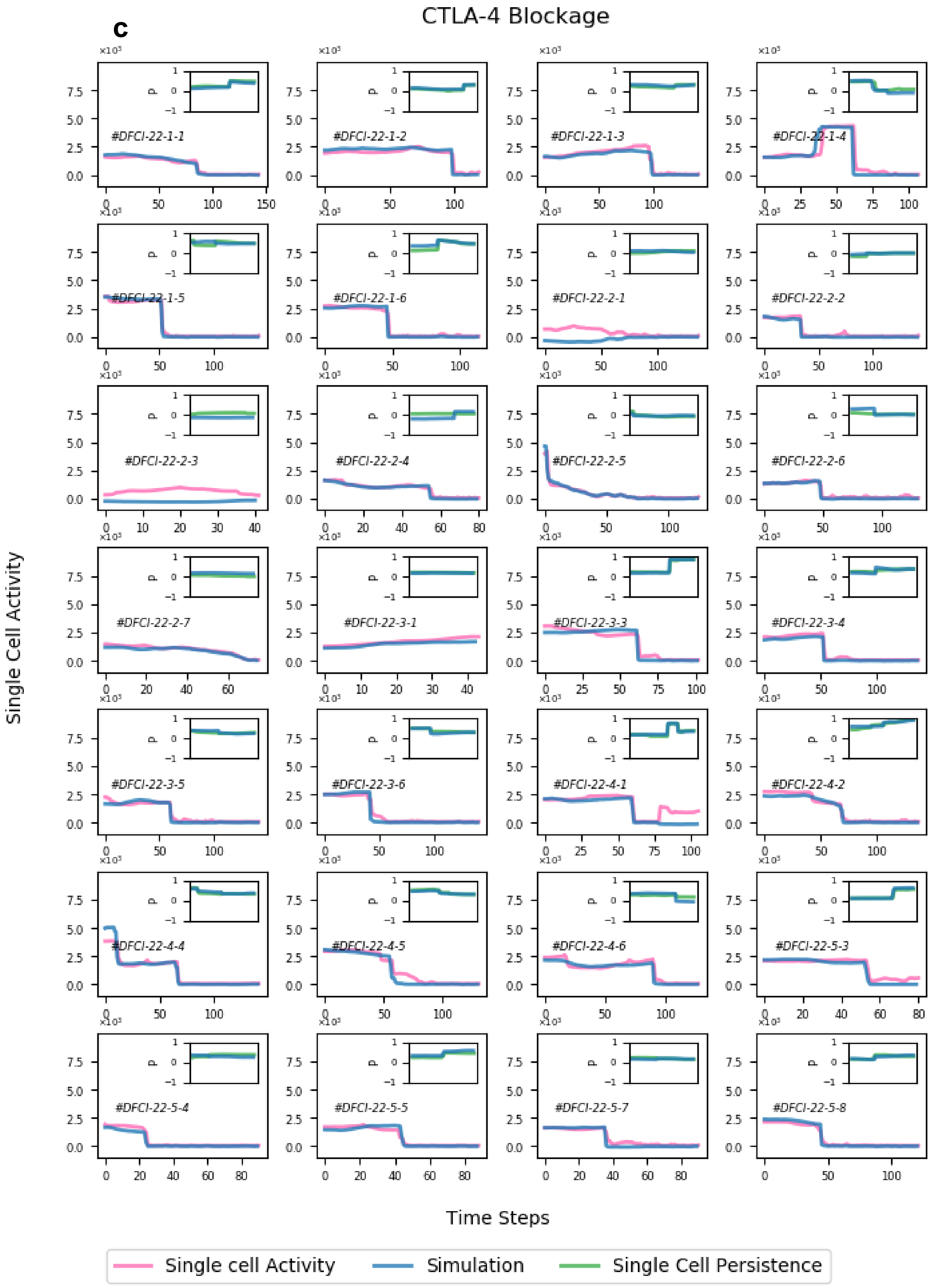
c
CTLA-4 Blockage
Single Cell Activity
Time Steps
#DFCI-22-1-1
#DFCI-22-1-2
#DFCI-22-1-3
#DFCI-22-1-4
#DFCI-22-1-5
#DFCI-22-1-6
#DFCI-22-2-1
#DFCI-22-2-2
#DFCI-22-2-3
#DFCI-22-2-4
#DFCI-22-2-5
#DFCI-22-2-6
#DFCI-22-2-7
#DFCI-22-3-1
#DFCI-22-3-3
#DFCI-22-3-4
#DFCI-22-3-5
#DFCI-22-3-6
#DFCI-22-4-1
#DFCI-22-4-2
#DFCI-22-4-4
#DFCI-22-4-5
#DFCI-22-4-6
#DFCI-22-5-3
#DFCI-22-5-4
#DFCI-22-5-5
#DFCI-22-5-7
#DFCI-22-5-8
Single cell Activity
Simulation
Single Cell Persistence

**Figure S2. Single cell parameters of AR-1 model of migration in combined inhibition of PD-1 and CTLA-4 pathways (a) as well as PD-1(b) and CTLA-4 blockage (c).** Bayesian inference method was applied on time series of cell trajectories and the mean values of the inferred activity parameter are shown for each cells (pink curves). The results of parameter estimation for AR1-simulated trajectories (blue curves) show good agreement with those derived from experiment. Insets: single cell persistence of the cells (dark green curves) and simulation-derived ones (light green curves). # notes indicate ID of patients from Dana Farber Cancer institute, the number of tumor spheroids and tracked cells number respectively.

#### Supplementary Note 3

#### Improving the initial values of lymphocyte subpopulations (correction of α and β parameters)

Considering that the initial values of activity for three subpopulations of lymphocytes (including PD-1-deficient lymphocytes, those partially expressing PD-1 and those perfectly possess this receptor) might have been unrealistic, the integrated model ran for a better estimation of $\alpha$ and $\beta$ parameters. In order to do so, using the already estimated $\alpha$ and $\beta$ parameters the model was simulated assuming again that the total activity initially belongs to the unexpressed subpopulation.

Model-derived distribution of CTLA-4 deactivated cells with different PD-1 expression states were compared. It was anticipated that the relative populations give a better estimation for the initial values of simulation which would result in a more accurate estimation of $\alpha$ and $\beta$ parameters under the condition of CTLA-4 inhibition.

However, the dominant proportion of lymphocytes population gathered in the stock of subpopulation lacking PD-1 receptor, and moreover, there was no improvement in the estimation of $\alpha$ and $\beta$ parameters when we used the obtained distribution of lymphocytes as initial values. Therefore, the presumption (model hypothesis) which initially assigns the total activity to the population lacking PD-1 receptor is an admissible assumption.
